# Population Structure and Antimicrobial Resistance Gene Transfer Respiratory *Escherichia coli* Isolated from Swine in China

**DOI:** 10.64898/2026.03.24.713904

**Authors:** Jianhai Li, Huiwen Mo, Chaofei Wang, Wenjian Cao, Jinchao Zhang, Shouneng Shi, Ruhua Qiu, Rui Fang, Junlong Zhao

## Abstract

Porcine respiratory diseases caused by extraintestinal pathogenic *Escherichia coli* (ExPEC) pose a severe threat to swine production and public health; however, research on respiratory tract-isolated ExPEC remains limited. This study comprehensively analyzed the genomic characteristics and antibiotic resistance gene (ARG) transfer potential of 441 ExPEC strains isolated from porcine lungs across 21 Chinese provinces (including 53 newly isolated strains from 2022–2024 and 388 NCBI-deposited strains). Phylogenetic analysis revealed that 84% of the isolates belonged to phylogroups A, B1, and C, with ST410, ST101, and ST88 as the predominant STs. The strains exhibited extensive ARG diversity, harboring 111 distinct ARG subtypes, with *sul2* (81.4%), *floR* (73.5%), and *tet (A)* (68.0%) being the most prevalent. Importantly, critical “last-resort” antibiotic resistance genes (e.g., *bla*_NDM-1/5_, the *mcr* family, and *tet (X4)*) were also detected. Notably, 77.2% of the ARGs presented horizontal transfer potential, with plasmids (especially IncF family replicons) serving as core vectors, followed by integrons and transposons. Cooccurrence network analysis identified *aph (3’’)-Ib*, *aph (6)-Id*, *sul2*, and *floR* as core subnetworks driving multidrug resistance dissemination. Pangenomic analysis confirmed an open genome architecture, with core genes accounting for only 6%, reflecting the strains’ capacity to acquire exogenous genetic material via horizontal transfer. From the One Health perspective, these transferable ARGs can spread to the environment and humans through fecal discharge and the food chain. These findings underscore the importance of monitoring and controlling ExPEC infections in swine, as such strains can as reservoirs of ARGs, pose potential risks to human health, and may even act as sources of pathogenic agents responsible for human infections.

**IMPORTANCE:** Porcine respiratory ExPEC-induced diseases threaten swine production and public health, yet respiratory tract-isolated ExPEC research remains scarce. This study comprehensively analyzed 441 porcine lung ExPEC strains across 21 Chinese provinces, uncovering their dominant phylogroups, high ARG diversity (111 subtypes) and the presence of “last-resort” antibiotic resistance genes. We identified 77.2% of ARGs with horizontal transfer potential, plasmids (especially IncF family) as core vectors, and a core ARG subnetwork driving multidrug resistance. The open pangenome (6% core genes) highlights ExPEC’s strong capacity to acquire exogenous genes. These findings fill the research gap of respiratory ExPEC, clarify ARG transmission mechanisms in swine ExPEC, and provide critical genomic data for One Health-based AMR surveillance and control, guiding targeted strategies to mitigate ARG spread from swine to humans and the environment.

## INTRODUCTION

Porcine respiratory diseases have a profoundly significant impact on global swine production, resulting in substantial economic losses characterized by impaired growth performance, increased treatment costs, and elevated mortality rates(1, 2). Porcine respiratory diseases are typically the consequence of the combined action of primary and opportunistic pathogenic agents(3). Previous studies have focused primarily on diseases such as porcine reproductive and respiratory syndrome virus (PRRSV), Streptococcus suis (SS), *Pasteurella multocida* (PM), and *Mycoplasma hyopneumoniae* (MHYO)(4, 5). Notably, in our clinical practice, extraintestinal pathogenic *Escherichia coli* (ExPEC) has been frequently isolated from the respiratory tract of pigs. Accumulating evidence has demonstrated that ExPEC may cause fatal pneumonia in swine, resulting in severe health threats to pigs(6). Although some studies have focused on porcine ExPEC strains in recent years(7, 8), research specifically addressing ExPEC isolates from the respiratory tract of pigs remains scarce.

To increase the health status and growth performance of swine, a variety of strategies have been implemented for the prophylaxis and therapy of swine colibacillosis, with antibiotics being the predominant intervention(9). Antimicrobial resistance (AMR) poses a substantial threat to global public health(10). In *Escherichia coli*, the evolution and dissemination of AMR are predominantly driven by antibiotic utilization, primarily via selective pressures that elicit the horizontal transfer of antibiotic resistance genes (ARGs) in this bacterium(11, 12). The extensive utilization of antibiotics in the Chinese swine industry has remarkably elevated the risk of ARG transfer(13). Studies have shown that the majority of *Escherichia coli* isolates from healthy and diseased swine as well as poultry in China are resistant to two or more antibiotics(14, 15).

Commensal and pathogenic bacteria originating from human, agricultural, and environmental settings all harbor ARGs within their genomes, with such genes amenable to horizontal transmission facilitated by mobile genetic elements(16). Plasmids, insertion sequences, transposons, integrative conjugative elements, and integrons constitute a repertoire of genomic structures that underpin the accelerated transmission of ARGs among diverse bacterial taxa(17). Commonly carrying both ARGs and virulence factors (VFs), antimicrobial-resistant *E. coli* relies on these genetic elements as key contributors to the development of drug resistance and the onset of clinical infections(18). Their inherent capacity for cross-host dissemination and horizontal transfer of such genetic elements facilitates the progressive accumulation of additional antimicrobial resistance attributes, a process that directly fuels the emergence of multidrug-resistant *E. coli* strains(19). Elucidating the transmission dynamics of ARGs among bacteria of human and animal host origins constitutes a core mandate of the One Health paradigm, and such insights can furnish critical evidence for formulating targeted and efficacious strategies to alleviate antimicrobial resistance(20, 21).

As the world’s largest swine-producing country, China inevitably relies on antibiotic administration during swine production. However, the inappropriate use of antibiotics has given rise to severe environmental and public health risks, with AMR representing a particularly prominent concern(22). Previous studies have demonstrated that extraintestinal pathogenic *Escherichia coli* (ExPEC) has the potential to cause substantial economic losses in the swine industry(23). Over the past decade, continuous advances in sequencing technologies have markedly improved the accessibility and throughput of whole-genome sequencing (WGS), thereby leading to the development of innovative strategies for the identification and in-depth characterization of pathogenic agents(24). Combined analysis of *E. coli* isolates derived from swine and other animal hosts via antimicrobial susceptibility testing and WGS has substantially advanced the understanding of the evolutionary dynamics and transmission pathways underlying AMR development and dissemination(25).

In the present study, we conducted a comprehensive analysis of the whole-genome characteristics and ARG transfer potential of 53 ExPEC isolates derived from porcine lungs in China between 2023 and 2024, as well as porcine lung-origin ExPEC strains deposited in the NCBI database. The primary objectives were to elucidate the genomic profiles of porcine lung-associated ExPEC in China and the current status of AMR in Chinese swine farms, which holds substantial significance for global AMR surveillance initiatives.

## MATERIALS AND METHODS

### ExPEC Isolate Collection

ExPEC isolates were collected from the lung tissues of pigs exhibiting various clinical signs of respiratory tract infections between 2022 and 2024 by the Animal Disease Diagnosis Laboratory of Wuhan Kevchunag Biology Technology Co. Ltd. Initially, pigs suspected of having respiratory tract infections were identified on the basis of clinical manifestations, including lateral recumbency and abdominal breathing. These clinical evaluations were conducted by trained veterinarians and animal health professionals. Lung tissues and other relevant organs were harvested from pigs that died of the aforementioned symptoms for further analysis. Tissue sampling was performed via sterile instruments, with operators wearing appropriate personal protective equipment. The samples were promptly collected, sealed in sterile containers, and stored under refrigeration to preserve their integrity. Detailed records were maintained regarding the sampling time and geographical location of the pig farms. The samples, together with the corresponding records, were transported to the Animal Disease Diagnosis Laboratory of Wuhan Kevchunag Biology Technology Co. Ltd. Upon receipt of the samples, the laboratory conducted further diagnostic tests to confirm the presence of ExPEC. The collected samples were predominantly sourced from Hubei Province and adjacent provinces.

### Short-read assembly and genome annotation

The genomes of all ExPEC isolates from porcine lungs were sequenced via the Illumina HiSeq 2500 platform. The raw short reads generated via the Illumina platform were initially quality-assessed with FastQC (https://www.bioinformatics.babraham.ac.uk/projects/fastqc/), followed by processing with Trimmomatic(26) to remove adapter sequences and low-quality reads. High-quality read sequences of each isolate were assembled via SPAdes(27). The completeness and contamination level of each assembled genome were evaluated with CheckM(28), and genomes with completeness < 95% or contamination > 5% were excluded from subsequent analyses. Finally, the draft genomes were functionally annotated via Bakta(29).

### Download of ExPEC genome data

Sequence files of an additional 388 ExPEC genomes isolated from porcine lung tissues were batch-downloaded from the National Center for Biotechnology Information (NCBI, https://www.ncbi.nlm.nih.gov/) via a custom script. All the genomes met the following quality control criteria: completeness > 90%, contamination rate < 5%, number of contigs ≤ 500, and N50 value ≥ 40,000.

### Bioinformatic analysis

We created a Python program (ARMobility) to identify ARGs and MGEs for bacterial nucleotide sequences via BLAST analysis. The transferability of ARGs is determined by evaluating the positional relationships between ARGs and MGEs(30). The identification of ARGs relies on the ResFinder database(31). Integrons were identified via the INTEGRALL database(32), transposable elements via the TRANSPOSON REGISTRY database(33), insertion sequences via the ISfinder database(34), and integrative and conjugative elements (ICEs) via the ICEberg database(35). Plispy is employed for plasmid sequence identification(36), whereas Phispy is utilized for prophage sequence recognition(37). PlasmidFinder was used to determine putative plasmids carried by the sequenced strains(38). In silico serogroups were determined by SerotypeFinder(39). In silico phylotyping was performed via Clermon typing(40). Identification of sequence types via pubMLST(41). A phylogenetic tree was constructed on the basis of core genome SNPs via the snippy-multipipeline integrated within the snippy. Phylogenetic tree inference was performed via FastTree(42), with the analysis based on core genome single nucleotide polymorphisms (SNPs) identified via the snippy-multipeline integrated within the snippy(43). Pangenome analysis was performed via Roary(44).

## RESULTS

### Serotype distribution, MLST identification, and differential clades of the isolates

The 53 ExPEC strains isolated from porcine lungs in this study were all obtained from clinically diseased swine collected between 2022 and 2024, with each strain derived from a different individual. The diseased tissue samples were collected from central China, mainly from Hubei Province (Figure 1 A). Phylogenetic analysis revealed that 47.2% of the isolates belonged to phylogroup B1 (n=25), 32.1% to phylogroup A (n=17), and 9.4% to phylogroup C (n=5). Phylogroup D and phylogroup G each comprised two isolates, whereas only one isolate was assigned to phylogroup B2 and phylogroup F, respectively (Figure 1 A). On the basis of in silico multilocus sequence typing (MLST), 51 of 53 ExPEC strains could be assigned to 25 STs, with 2 strains representing novel STs. ST11 was the predominant type (n=7), followed by ST162 (n=5) and ST641 (n=5), while the remaining STs were detected at a low frequency (n<5) (Figure 1 A).

**Fig 1.**
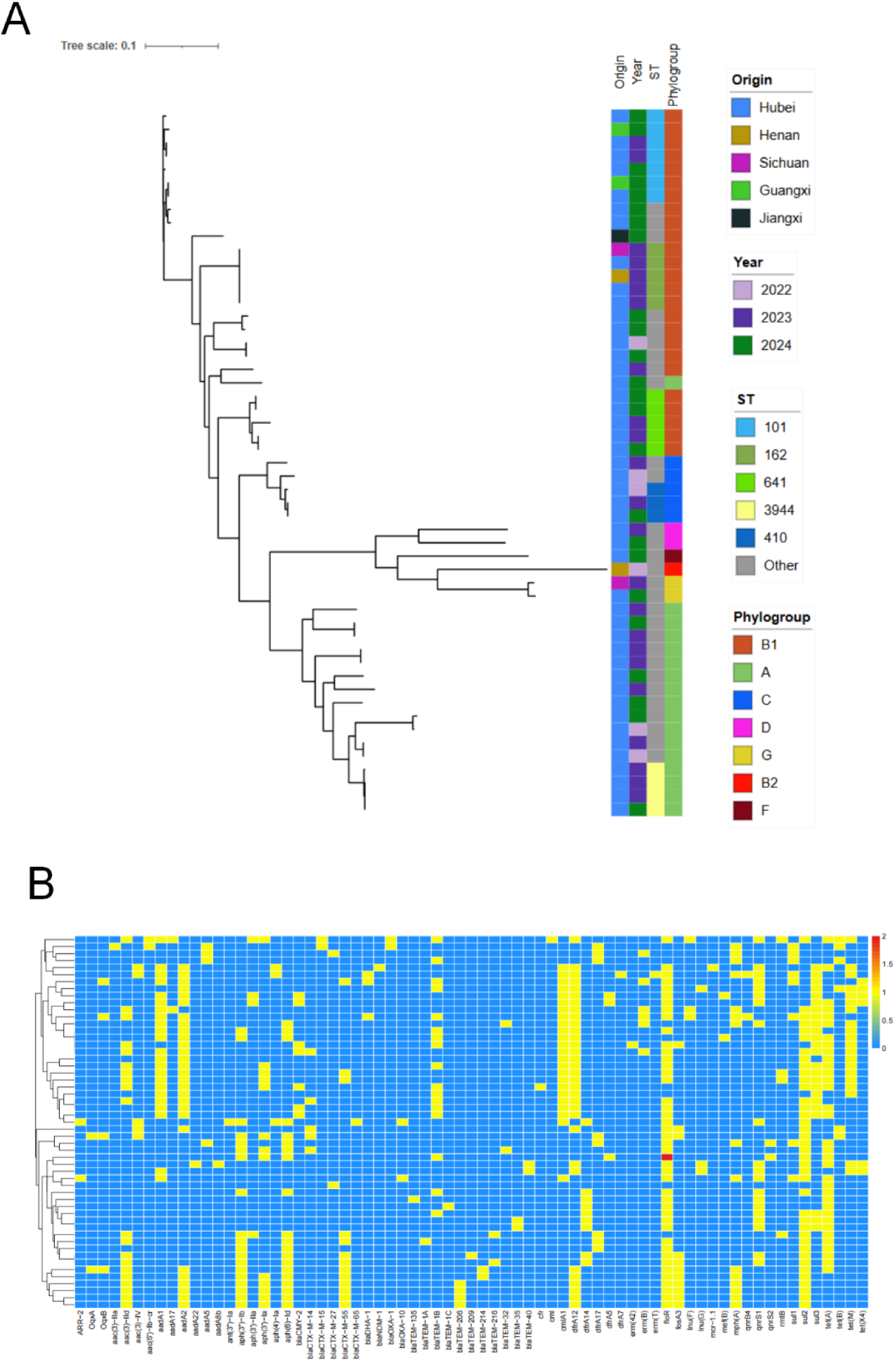
The maximum likelihood phylogenetic tree and the deletion/presence of ARGs in ExPEC isolated from pig respiratory tracts during 2022 to 2024. **A** Maximum likelihood phylogenetic tree of 53 swine-derived ExPEC isolates. From left to right, the four columns represent Isolation Province, Isolation Date, ST Type and Phylogroup respectively. **B** The carriage status of antimicrobial resistance genes in 53 porcine-derived extraintestinal pathogenic Escherichia coli (ExPEC) isolates.

### Resistome repertoires of ExPEC isolates

The 53 ExPEC isolates presented high prevalence and functional convergence in terms of ARGs. A total of 69 subtypes of ARGs were detected in 53 ExPEC isolates. The core highly prevalent resistance genes and their corresponding prevalence rates are listed as follows: *floR* (84.9%), *sul2* (73.6%), *tet(A)* (64.2%), *dfrA12* (60.4%), *aadA2* (52.8%), *cmlA1* (45.3%), *sul3* (45.3%), *aadA1* (43.4%), *aac*(*3*)*-IId* (37.8%), *aph(6)-Id* (37.8%), *qnrS1* (37.8%) *tet(M)* (37.8%), *aph(3’’)-Ib* (35.8%), and *bla*_TEM-1B_ (32.1%) (Figure 1 B). Among the detected ARGs, several are highly prevalent. A high overall prevalence of ARGs was observed in the isolates. The highly prevalent ARGs identified in this study cover a wide spectrum of antimicrobial classes, including aminoglycosides, sulfonamides, tetracyclines, trimethoprim, florfenicol, chloramphenicol, quinolones and β-lactams. The high prevalence of these antibiotics is likely to induce therapeutic failure of the aforementioned antibiotics against ExPEC, thereby increasing the difficulty and risk of treatment failure in patients with clinical infections. Moreover, these highly prevalent resistance genes tend to spread among strains via horizontal gene transfer, accelerating the emergence and dissemination of multidrug-resistant isolates and further intensifying the selective pressure on clinical antimicrobial agents.

### Phylogenomic analysis of ExPEC isolates from swine lungs in China

As of June 2025, we screened the NCBI database for entries with complete isolation metadata from swine lungs in China. After quality control filtering (removing genomes with completeness <95% and contamination >5%), a total of 388 high-quality ExPEC genomes from Chinese swine lungs were obtained. Combining these results with those of the 53 isolates obtained in the present study, a comprehensive analysis was performed on a total of 441 extraintestinal pathogenic ExPEC isolates derived from swine lungs in China. In terms of geographical distribution, these genomes were distributed across 21 provinces or municipalities directly under the Central Government of China. The majority of them originated from Central China, with Hubei Province accounting for the highest proportion (46.7%), followed by Henan, Hunan, and Zhejiang Provinces at 11.1%, 8.6%, and 7.0%, respectively (Figure 2 A). Phylogenetic analysis revealed that 30.4% of the isolates belonged to phylogroup A, 30.2% to phylogroup B1, 23.6% to phylogroup C, 6.1% to phylogroup F, and 5.7% to the recently defined phylogroup G. Notably, the ExPEC isolates investigated herein were rarely assigned to phylogroups B2, D, and E, with the proportion of each phylogroup accounting for less than 3% (Figure 2 B). ExPEC strains isolated from the lungs of swine in China exhibit high diversity. On the basis of in silico MLST, 424 of the 441 ExPEC strains could be assigned to 106 STs, with 17 strains representing novel STs. The top 5 STs with the most isolates included ST410 (13.8%), ST101 (7.3%), ST88 (6.3%), ST117 (5.0%) and ST156 (4.8%), accounting for more than 37% of the entire collection. Among these STs, ST410 and ST88 belonged to phylogroup C, ST101 and ST156 belonged to phylogroup B1, and ST117 was categorized into phylogroup G (Figure 2 B). Our findings demonstrated that strains isolated at specific time points are distributed throughout the entire phylogenetic tree, indicating that there is no association between isolation time and phylogroups or STs. We quantified the number of ARGs and VFs harbored by each bacterial strain across different phylogenetic groups. Compared with the other phylogenetic groups, the B2 group presented a lower abundance of ARGs, while its VF content was significantly greater. Notably, the phylogenetically prevalent groups A, B1, and C presented a greater number of ARGs but a lower count of VFs than did the other groups did (Figure 2 C).

**Fig 2.**
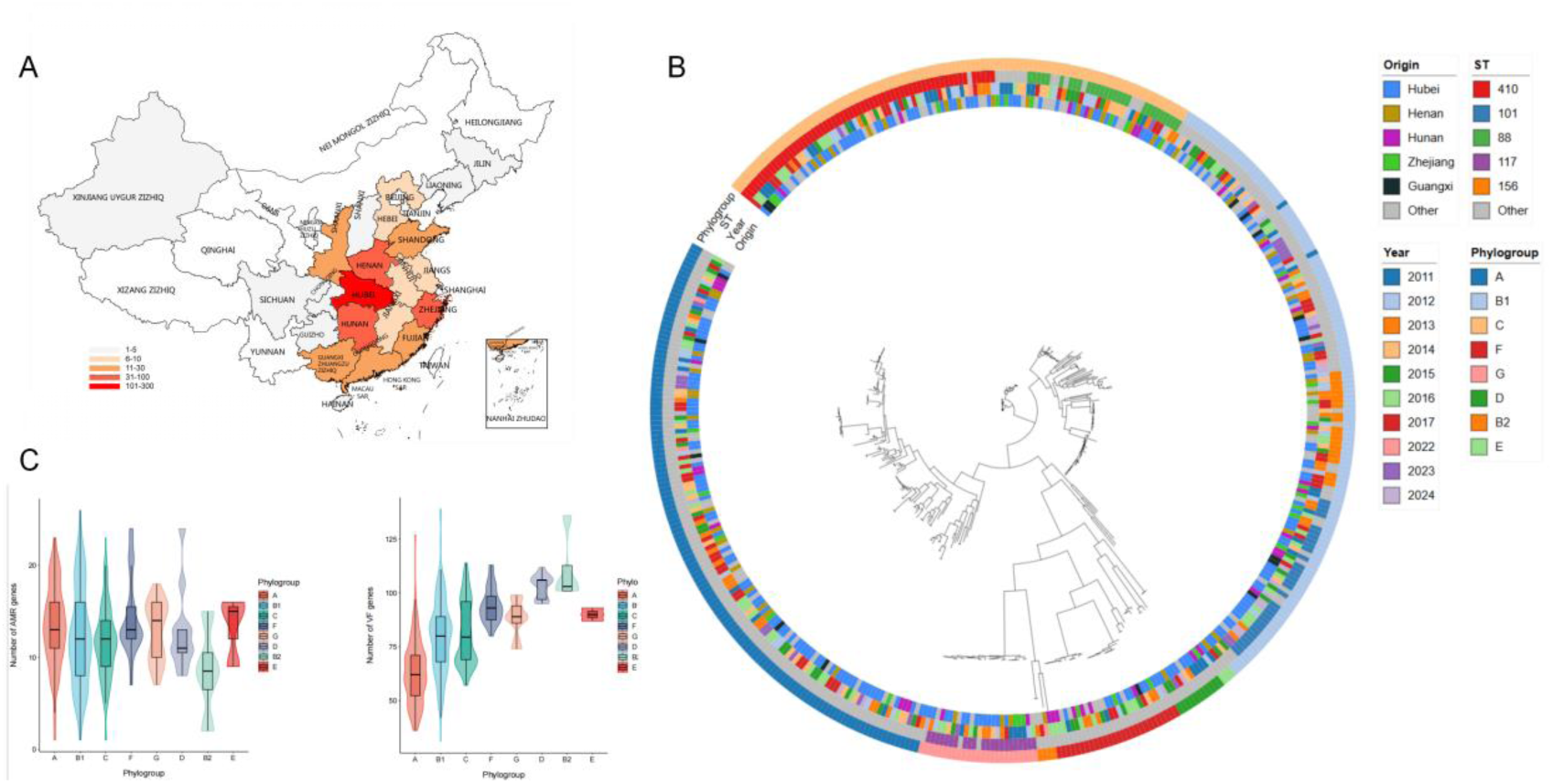
The isolation sources and genomic characteristics of 441 ExPEC isolates from swine respiratory tracts. **A** China map showing the number of porcine ExPEC isolates from pig farm lungs in each province of China. **B** Maximum likelihood phylogenetic tree of 441 swine-derived ExPEC isolates. From inner to outer, the four columns represent Isolation Province, Isolation Date, ST Type and Phylogroup respectively. **C** Statistics of the number of ARGs and VFs in ExPEC isolates across different phylogroups.

### Resistome repertoire of ExPEC isolates from swine lungs in China

The results of ARG detection in 441 ExPEC strains isolated from swine lungs revealed the identification of 111 distinct ARG subtypes, indicating a high carriage rate and remarkable diversity of ARGs among the tested strains. Notably, the genomes of 31 ExPEC strains harbored at least two copies of *sul2*, 21 genomes carried no fewer than two copies of *bla*_TEM-1B_, and the numbers of genomes containing at least two copies of *sul1* and *tet(A)* were 16 and 14, respectively (Supplementary Data 1). A statistical analysis of the prevalence of all resistance genes revealed that the sulfonamide resistance gene *sul2* was the most prevalent gene, with 81.4% (n=359) of the isolates harboring this resistance gene. This was followed by the chloramphenicol resistance gene *floR* (n=324, 73.5%), the tetracycline resistance gene *tet(A)* (n=300, 68.0%), and the β-lactam resistance gene *bla*_TEM-1B_ (n=282, 63.9%). Among the aminoglycoside resistance genes, *aph(6)-Id* (n=274, 62.1%) and *aph(3’’)-Ib* (n=272, 61.7%) had the highest detection rates. Additionally, the *aadA* gene family (including *aadA2* with n=162 and *aadA1* with n=136) was also widely prevalent, indicating that aminoglycoside resistance is relatively common in this bacterial population. The β-lactam resistance gene family is highly diverse. In addition to the dominant gene *bla*_TEM-1B_, 12 other *bla*_TEM_ subtypes (including *bla*_TEM-1A_ and *bla*_TEM-206_) and 14 *bla*_CTX-M_ subtypes (such as *bla*_CTX-M-14_ (n=76), *bla*_CTX-M-55_ (n=64), and *bla*_CTX-M-65_ (n=26)) were detected. Moreover, ARGs conferring resistance to cephalosporins and penicillins, including *bla*_CMY-2_ (n=38) and *bla*_OXA-1_ (n=39), were also identified. More notably, the carbapenem resistance genes *bla*_NDM-1_ (n=2) and *bla*_NDM-5_ (n=3) were both detected.

Although their prevalence rates are relatively low (0.45%∼0.68%), they pose a serious threat to clinical treatment because of the carbapenem resistance mediated by these genes. In addition, resistance genes corresponding to important “last-resort” antibiotics were detected: a total of 7 subtypes of the colistin resistance gene *mcr* family were identified, among which *mcr-1.26* (n=50, 11.3%) and *mcr-1.1* (n=38, 8.6%) were the predominant prevalent subtypes. The tigecycline resistance gene *tet(X4)* was detected 7 times (1.6%), whereas the fosfomycin resistance gene *fosA3* (n=73, 16.6%), quinolone resistance gene *qnrS1* (n=69, 15.6%), and high-level aminoglycoside resistance gene *rmtB* (n=68, 15.4%) presented relatively high prevalence rates. These findings further indicate the resistance risk of *E. coli* in this population to multiple classes of critical antibiotics (Figure 3 A) (Supplementary Data 2).

**Fig 3.**
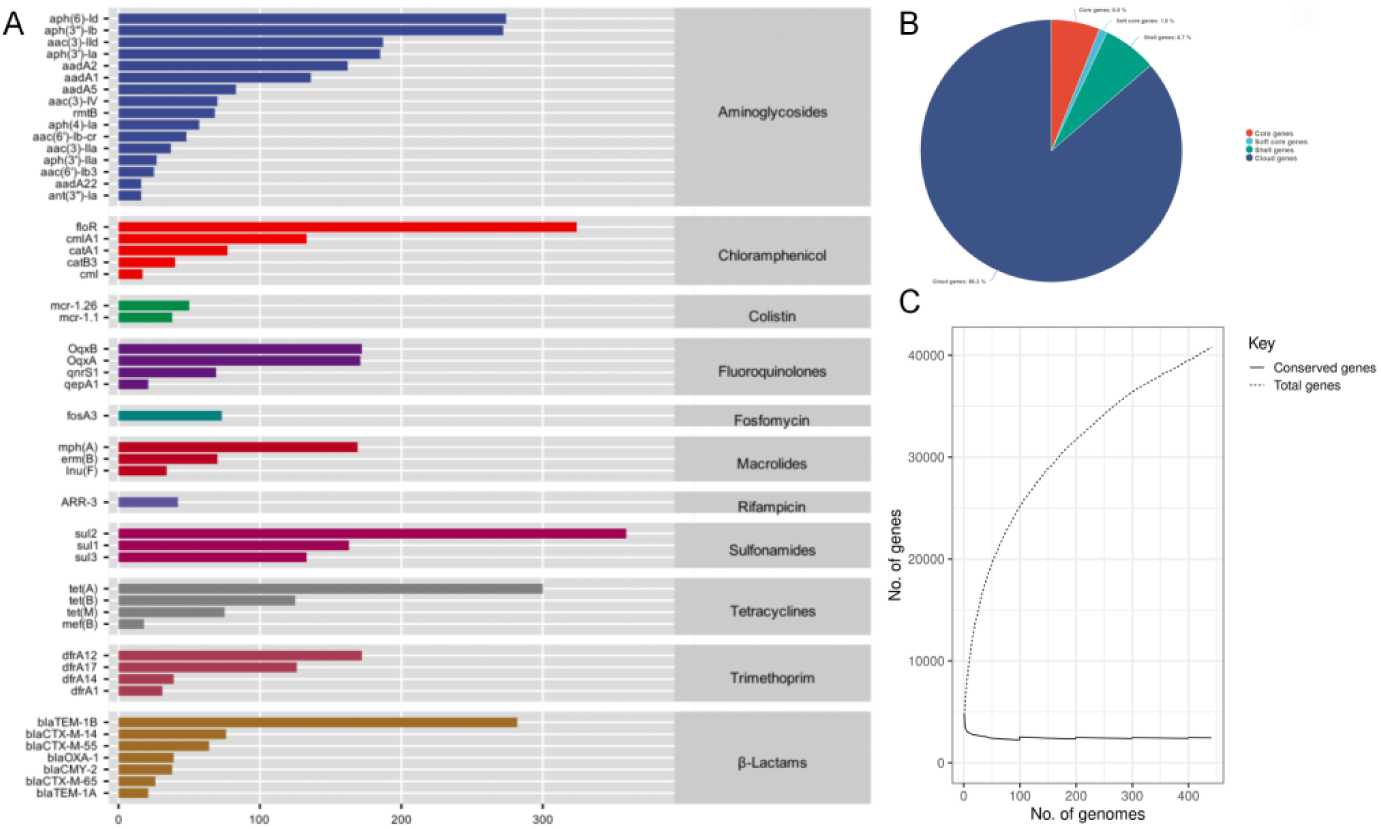
The prevalence of ARGs and pan-genome characteristics of 441 ExPEC strains. **A** The bar chart illustrates the number of each antimicrobial resistance gene detected in 441 ExPEC isolates. Colors represent the classes of antimicrobial agents to which resistance is conferred by the genetic determinants. This figure only displays representative high-prevalence ARGs; complete information is available in the supplementary data. **B** Distribution of gene frequencies across the 441 ExPEC genomes. **C** Gene accumulation curves reveal pangenome expansion and a nearly constant core genome with increasing genome numbers.

To investigate the genetic determinants behind MDR (extensive drug resistance, nonsusceptibility to at least one agent in all but two or fewer antimicrobial categories) *E. coli*, we examined the compositional profiles of ARGs in each genome. In the 436 genomes containing at least 2 antibiotic resistance genes (ARGs), a total of 389 distinct ARG combinations were identified. The results revealed that 80.5% (n=351) of the genomes harbored a unique set of ARG combinations. Each combination contained an average of 13 ARGs, with the largest one containing up to 25. Among the 389 identified ARG combinations, 76.3% contained no fewer than 10 ARGs per combination, whereas 8.5% contained 20 or more ARGs per combination (Supplementary Data 3). The results of ARG combination revealed a remarkable diversity of ARG combinations in the majority of ExPEC genomes isolated from swine lungs.

### Genome characteristics of ExPEC isolates from swine lungs in China

For the 441 ExPEC strains isolated from the lungs of Chinese swine, the average genome length was 4,435,718.2 bp, with an average GC content of 51.67%. Each genome contained an average of 4,841.4 CDSs, 8.4 rRNAs, and 147.6 tRNAs (Table 1). The results from the pangenomic analysis of 441 *Escherichia coli* strains revealed that only 6% of the genes were core genes, whereas 86.3% were cloud genes (Figure 3 B). This finding indicates remarkable genomic plasticity in *E. coli* strains isolated from the lungs. Further pangenomic analysis revealed that ExPEC strains isolated from porcine lungs exhibit an open pangenome architecture. As the number of genomes included in the analysis increased, the size of the gene cluster pool continued to expand, indicating considerable genetic diversity among these porcine lung-isolated ExPEC strains. In contrast, the core genome displayed a declining trend, suggesting gene loss with the addition of more genomes (Figure 3 C). This plasticity implies that extraintestinal pathogenic ExPEC strains derived from porcine lungs can continuously acquire exogenous genetic materials — including antibiotic resistance genes and virulence genes — through mechanisms such as horizontal gene transfer (HGT).

**Table 1.** presents the genomic information of 441 ExPEC isolates.

| Genome length | GC content | CDS | rRNA | tRNA |
| --- | --- | --- | --- | --- |
| 4,435,718.2 bp | 51.67% | 4,841.4 | 8.4 | 147.6 |

### ARG transfer analysis of ExPEC isolates from swine lungs in China

We utilized a custom-developed Python program to analyze the transfer potential of all resistance genes. The criteria for determining horizontally transferable ARGs are as follows: 1) For an ARG, if the same IS sequence is present on both sides within 10 kb upstream and downstream of the gene, the ARG is considered to possess transfer potential; 2) if an ARG is located within the sequence boundaries of IN, Tn, plasmids, prophage or ICE, it is deemed to have the potential for horizontal transfer. A total of 5,806 ARGs were detected in 441 strains of ExPEC, among which 77.2% (n=4,482) exhibited horizontal transferability (Figure 4 A) (Supplementary Data 4). Further analysis revealed that trimethoprim resistance genes (such as *dfrA12* and *dfrA17)* presented an exceptionally high average transferability rate of 97.82%, with *dfrA12* and *dfrA1* reaching 100%, representing the category with the highest risk of horizontal transmission. Chloramphenicol resistance genes (such as *floR*, *cmlA1*) and macrolide resistance genes (such as *mph(A)*, *erm(B)*) demonstrated remarkably strong overall transfer potential, with average transferability rates of 89.03% and 88.06%, respectively, and over half of these genes presented a transferability rate ≥ 90%. Aminoglycoside resistance genes (such as *aph* series and *aad* series) had an average transferability rate of 86.3%. For quinolone resistance genes (such as *OqxB* and *qnrS1*), tetracycline resistance genes (*tet(A)*, *tet(B)*, and *tet(M)*), and sulfonamide resistance genes (*sul2*, *sul1*, and *sul3*), the average transferability rates ranged between 70% and 85%. Notably, tetracycline- and quinolone resistance genes presented significant intracategory variability in transferability (e.g., *tet(M)* presented 100% transferability, whereas *tet(B)* presented only 55.20%), whereas all sulfonamide resistance genes presented moderate transfer characteristics. β-lactam resistance genes (such as the *bla*_TEM_ series and *bla*_CTX-M_ series) had an average transferability rate of 66.49%, with the highest intergenic heterogeneity, encompassing both high-transfer genes such as *bla*_OXA-1_ (100%) and low-transfer genes such as *bla*_CMY-2_ (18.42%). The average transferability rate of colistin resistance genes (*mcr-1.26*, *mcr-1.1*) was 50.56%, indicating a relatively lower risk of horizontal transmission than other resistance genes (Figure 4 B) (Supplementary Data 4). Additionally, one representative gene each from the rifampicin-resistant (*ARR-3*) and fosfomycin-resistant (*fosA3*) categories presented transferability rates of 100% and 84.93%, respectively. These results reflect the diverse transfer characteristics of resistance genes across different antibiotic classes.

**Fig 4.**
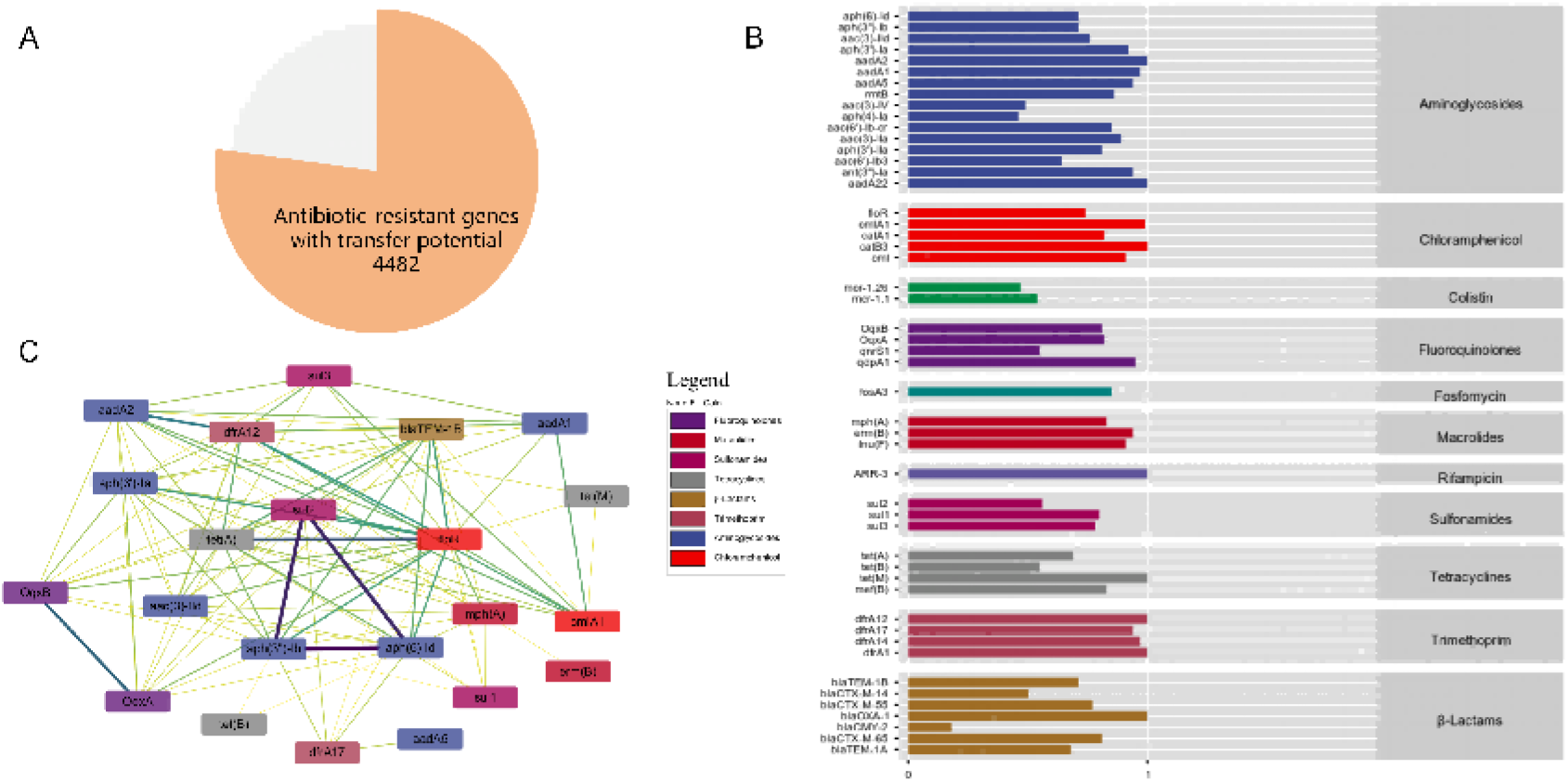
Diversity of genetic determinants of antibiotic resistance in 441 ExPEC strains. **A** The pie chart shows the proportion of transferable ARGs in all detected ARGs. **B** The bar chart presents the proportion of each ARG with transfer potential detected in 441 ExPEC isolates. Different colors represent the classes of antimicrobial agents to which these genetic determinants confer resistance. This figure only displays representative high-prevalence ARGs; complete information is available in the supplementary data. **C** Co-occurrence network of antimicrobial resistance genes (ARGs) in ExPEC strains. The network was constructed based on the co-occurrence frequency of pairwise ARGs with transfer potential across 441 genomes. The thickness and color intensity of each connection (edge) between ARGs (nodes) correspond to the frequency of co-occurrence of the two genes in the same genome. This figure only displays genes that co-occur in at least 50 genomes. All connection information is available in Supplementary Data.

To investigate the simultaneous transfer of multiple ARGs, we conducted co-occurrence network analysis on the entire set of ARGs. Cooccurrence network analysis revealed that *aph(3’’)-Ib*, *aph(6)-Id*, *sul2*, and *floR* constitute the most tightly connected core subnetwork, with the majority of other ARGs linked to this subnetwork. Within this core subnetwork, the co-occurrence frequency between *aph(3’’)-Ib* and *aph(6)-Id* was the highest (n=185), followed by *aph(3’’)-Ib* with *sul2* (n=170) and *aph(6)-Id* with *sul2* (n=168). For *floR*, it cooccurs with *sul2* 108 times and with both *aph(3’’)-Ib* and *aph(6)-Id* 104 times each. Notably, *aph(3’’)-Ib* and *aph(6)-Id* are both aminoglycoside resistance genes, *sul2* mediates sulfonamide resistance, and *floR* confers chloramphenicol resistance. The synergistic co-occurrence of these four genes directly facilitates the dissemination of the “aminoglycoside-sulfonamide-chloramphenicol” multidrug-resistant phenotype. Furthermore, this core subnetwork forms extended associations with *tet(A)* (a tetracycline resistance gene cooccurring with *floR* 149 times) and *bla*_TEM-1B_ (a β-lactam resistance gene cooccurring with *floR* 106 times), among other genes, which further expands the coverage of the resistance spectrum (Figure 4 C) (Supplementary Data 5).

Beyond the core subnetwork, multiple high-frequency associated gene pairs were still identified in the co-occurrence network, reflecting the diverse characteristics of ARG co-occurrence. Among these genes, *OqxA* and *OqxB* had the highest co-occurrence frequency (n=140); both are efflux pump-related genes that synergistically mediate resistance to quinolones and other drugs. The gene pairs *aadA2* and *dfrA12* cooccurred 130 times, encoding aminoglycoside- and trimethoprim resistance-related proteins, respectively. Other important high-frequency co-occurrence combinations included *cmlA1* and *dfrA12* (n=101), *bla*_TEM-1B_ and *tet(A)* (n=99), and *aph(6)-Id* and *bla*_TEM-1B_ (n=98). The results of the mobile ARG co-occurrence network analysis indicated that the dissemination of mobile ARGs is not random. Instead, it centers on *aph(3’’)-Ib*, *aph(6)-Id*, *sul2*, and *floR* as the core, accompanied by the synergistic diffusion and transmission of multiple types of ARGs (Figure 4 C) (Supplementary Data 5).

### Correlation analysis between transferable ARGs and MGEs

To decipher the vectors of ARGs, we analyzed the associations between ARGs and MGEs. The results demonstrated that the number of ARGs associated with each MGE varied dramatically. Among these MGEs, plasmids presented the highest ARG carriage capacity, with 3,757 associated ARGs, which was substantially greater than the counts detected in integrons (n=1109), transposons (n=499), insertion sequences (n=94), and ICEs (n=76). In contrast, only 5 ARGs were linked to prophages. With respect to gene distribution, the vast majority of these ARGs (n=3,462) were MGE specific. Specifically, plasmid-specific ARGs predominated, accounting for 2,767 ARGs, followed by transposon-specific (n=388) and integron-specific (n=210) ARGs; notably, no phage-specific ARGs were identified. In addition, a total of 982 ARGs were shared across two or more MGEs. Among these shared ARGs, integrons and transposons represented a relatively large number (n=23), whereas plasmids represented varying numbers of ARGs with ISs and prophages. No shared ARGs were detected between certain MGE pairs, and only a small subset of ARGs were concurrently associated with three or more MGE types. In summary, the association profile of transferable ARGs with MGEs is characterized by three key features: plasmids serve as the core carriers, most ARGs are MGE specific, and a small proportion of ARGs are shared across multiple MGE types (Figure 5 A).

**Fig 5.**
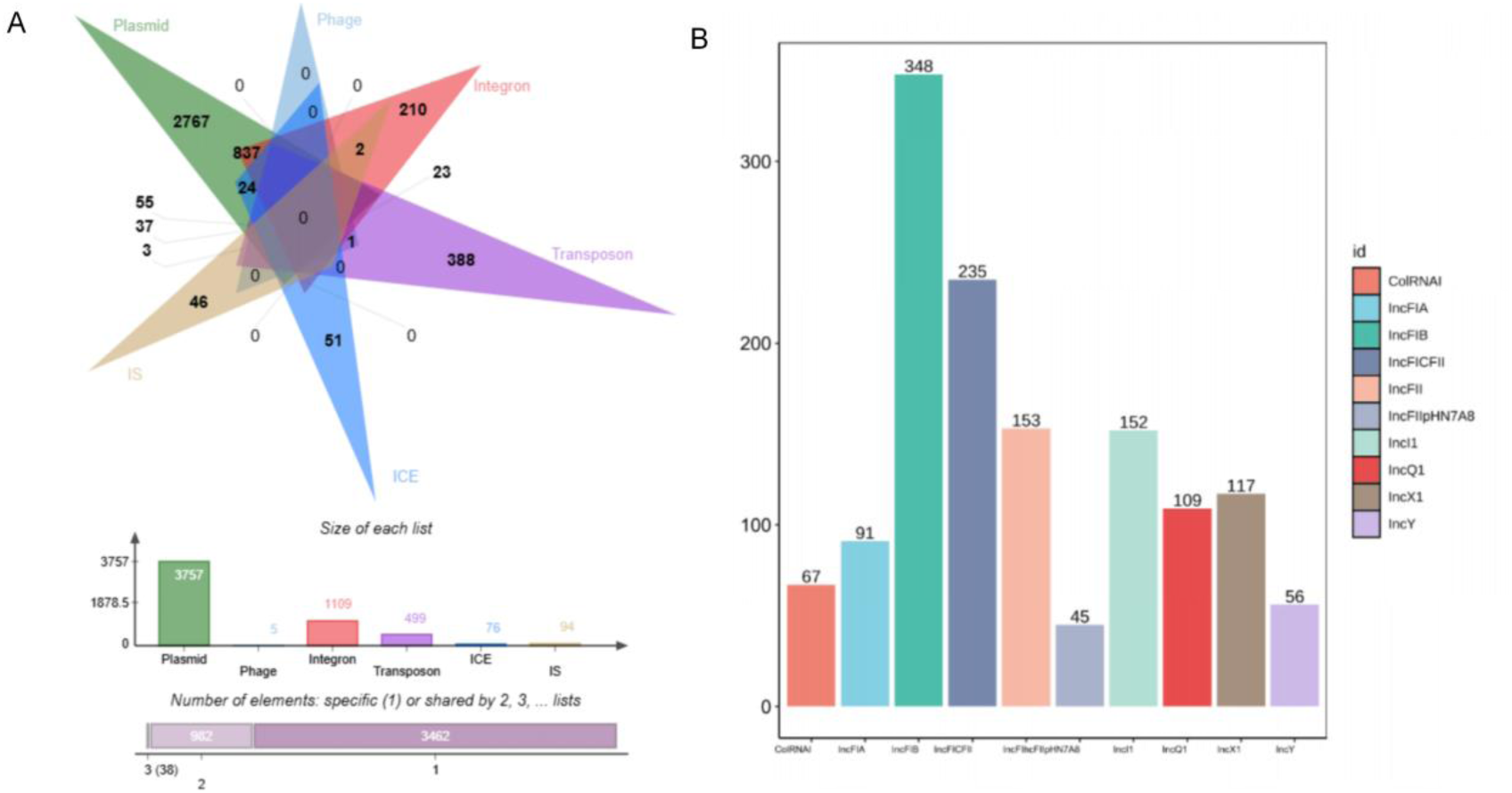
MGE and Plasmid Replicon Types in 441 ExPEC. **A** The Venn diagram illustrates the relationship between MGEs and ARGs with transfer potential. **B** The bar chart shows the number of distinct plasmid replicon types detected in 441 ExPEC isolates. Only representative high-prevalence plasmid replicons are displayed in this figure.

We subsequently detected the plasmid replicons of all ExPEC strains, with a total of 43 distinct types identified. Among these replicons, IncFIB (AP001918) was the predominant type, accounting for the highest proportion (348/1906) of all detected replicons. This was followed by IncFIC (FII) (235/1906) and IncFII (153/1906), with the combined proportion of these three replicons reaching 38.6%. Collectively, these genes constitute the core dominant group of plasmid replicons in this ExPEC population. Notably, replicons associated with the IncF family (including IncFIB, IncFIC, IncFII, and IncFIA) exhibited an overall extremely high detection frequency, which underscores the epidemic predominance of IncF-type plasmids in ExPEC. Typically, harboring multiple ARGs and VFs, these replicons serve as key vectors mediating the horizontal transmission of ARGs. In addition, the IncI1 and IncX1 replicons were detected 152 and 117 times, respectively, whereas the IncQ1 and IncFIA replicons were identified 109 and 91 times, respectively (Figure 5 B). These replicons also represent common plasmid replicon types in the studied population.

## DISCUSSION

We performed a genomic analysis of 441 ExPEC strains isolated from porcine lungs in China, and the results revealed that approximately 84% of the isolates belonged to phylogroups A, B1, and C. Compared with *E. coli* strains isolated from porcine feces, the ExPEC isolates derived from porcine lungs presented a significantly greater proportion of phylogroup C(45). The high-frequency STs, including ST410, ST101, ST88, ST117 and ST156, are consistent with the findings of a previous study focusing on Chinese porcine ExPEC(8). Notably, ST156 was identified as one of the predominant ST types in a study involving the isolation of *Escherichia coli* from retail meat across 22 cities in China(46). ExPEC strains carried by pork can lead to intestinal colonization and subsequent secondary urinary tract infections(47). These findings indicate that the potential transmission of extraintestinal pathogenic ExPEC between humans and swine may constitute a major source of human ExPEC infections. The results of antimicrobial resistance gene detection revealed that the sulfonamide resistance gene *sul2* (81.4%), chloramphenicol resistance gene *floR* (73.5%), and tetracycline resistance gene *tet(A)* (68.0%) were the most prevalent resistance determinants. This finding is highly consistent with historical medication practices in the swine industry of China, where these three classes of antibiotics are extensively administered for the prevention and treatment of porcine respiratory tract infections(48). This finding confirms that the abuse of antibiotics is one of the core driving factors underlying the prevalence of ARGs(49). Moreover, this study simultaneously detected carbapenem resistance genes (*bla*_NDM-1/5_), colistin resistance genes of the *mcr* family (seven subtypes), and the tigecycline resistance gene *tet(X4)* in porcine lung-derived ExPEC. Despite the relatively low prevalence of some of these genes (e.g., *tet(X4)* at 1.6%), the resistance mediated by such genes is associated with a near-total lack of effective therapeutic agents(50). These genes possess transmission potential and pose a severe threat to public health(51). Furthermore, a total of 111 antimicrobial resistance gene subtypes were identified across 441 bacterial isolates, with 80.5% of these strains harboring unique resistance gene combinations. Each isolate carried an average of 13 resistance genes, with the maximum number reaching 25. These findings indicate that porcine lung-derived ExPEC has evolved into a complex reservoir of antimicrobial resistance genes. The widespread prevalence of multidrug-resistant (MDR) and even extensively drug-resistant (XDR) phenotypes severely restricts options for clinical treatment regimens and further exacerbates economic losses in the swine industry.

The present study revealed that 77.2% of antibiotic resistance genes (ARGs) presented the potential for horizontal transfer, with plasmids serving as the predominant transmission vector (associated with 3,757 ARGs). Among these plasmids, the IncF replicon family (including IncFIB, IncFIC, IncFII, etc.) represented the dominant replicons, accounting for 52.7% of the total. IncF-type plasmids typically harbor multiple ARGs and VFs and possess a strong conjugative transfer capability; their prevalence advantage directly accelerates the dissemination of antibiotic resistance genes across different bacterial strains(52, 53). In addition, 152 IncI1 replicons and 117 IncX1 replicons were detected. Both the IncI1 and IncX1 plasmids harbor complete conjugative transfer elements(54), among which the IncX1 plasmid also exhibits a highly efficient conjugative transfer capability that enables rapid transmission across different bacterial strains and even distinct species(55). The presence of these two types of plasmids with strong transfer potential indicates that antibiotic resistance genes can spread to other pathogenic Enterobacteriaceae via horizontal gene transfer (HGT), which further increases the prevalence of resistant strains and accelerates the emergence of MDR strains.

Horizontal transfer of antibiotic resistance genes is one of the key factors driving the emergence of multidrug-resistant bacteria(16). Network analysis of cooccurring transferable ARGs revealed that *aph(3’’)-Ib*, *aph*(*6*)*-Id*, *sul2* and *floR* constitute a core associated subnetwork that synergistically mediates the dissemination of the aminoglycoside-sulfonamide-chloramphenicol multidrug resistance phenotype. Moreover, this subnetwork forms extended associations with genes such as *tet(A)* and *bla*_TEM-1B_, thereby further expanding the coverage of the resistance spectrum. This finding contributes to a better understanding of the mechanisms underlying the development of the multidrug-resistant phenotype in ExPEC. In addition, the high transfer rates of trimethoprim resistance genes (97.82%), chloramphenicol resistance genes (89.03%) and macrolide resistance genes (88.06%) indicate that such resistance genes can rapidly spread via MGEs. Moreover, plasmids, integrons and transposons share a portion of resistance genes, forming a pattern of synergistic transmission across multiple vectors, which further complicates the prevention and control of antimicrobial resistance genes(56–58). From the perspective of One Health, the transferable resistance genes carried by porcine lung-derived ExPEC can disseminate to the environment and humans through pathways such as fecal discharge and the food chain(59), thereby serving as critical intermediate nodes in the human-animal-environment antimicrobial resistance transmission chain.

Compared with previous studies, this research is the first to focus on porcine respiratory-specific ExPEC isolates, filling the research gap dedicated to respiratory ExPEC and addressing the limitation that prior studies have predominantly concentrated on intestinal isolates. Although the data are mainly concentrated in the central provinces of China, with insufficient coverage of the eastern, western, and northern regions, which may introduce regional biases in the analysis of population structure and antimicrobial resistance profiles, the dataset covers several major pig-producing provinces in China. Through a systematic pangenomic analysis of porcine lung-derived ExPEC isolates from 21 provinces in China, this study confirmed that the lung-derived ExPEC harbors an open pangenome, with core genes accounting for only 6% of the total gene repertoire. This study reveals the evolutionary strategy by which strains continuously acquire exogenous genetic material via horizontal gene transfer. Furthermore, this study quantifies the disparities in transfer potential among different classes of ARGs and identifies the core co-occurrence gene network as well as the key transmission vectors.

## Acknowledgments

This work was funded by the authors. We would like to acknowledge the support of Wuhan Kevchunag Biology Technology Co., Ltd., for this study.

## Conflict of Interest Statement

The authors declare no conflict of interest. Wuhan Kevchunag Biology Technology Co., Ltd. provided technical support for sample collection in this study with no other involvement in the research.

## AUTHOR CONTRIBUTIONS

Jianhai Li, Formal analysis, Investigation, Writing – original draft, Visualization |Huiwen Mo, Methodology, Software, Data curation |Chaofei Wang, Investigation, Resources, Validation |Wenjian Cao, Formal analysis, Methodology, Software |Jinchao Zhang, Resources, Data curation, Investigation |Shouneng Shi, Formal analysis, Visualization, Writing – review and editing |Ruhua Qiu, Resources, Supervision, Validation |Rui Fang, Methodology, Project administration, Writing – review and editing |Junlong Zhao, Conceptualization, Funding acquisition, Project administration, Supervision, Writing – review and editing.

## Ethics Statement

All animal manipulation procedures complied with the animal welfare regulations of the China Experimental Animal Center and the ethical standards of the Animal Ethics Committee of Huazhong Agricultural University (HZAUSW-2022-0007, HZAUSW-2022-0008). Samples were collected from naturally deceased pigs with written informed consent from farm owners, and no live animal experiments were conducted. All bacterial experiments followed institutional biosafety protocols.

## DATA AVAILABILITY

Sequencing data have been deposited in NCBI BioProject under accession number PRJNA1437841 (data will be made publicly available upon acceptance).

